# A microtubule associated protein is essential for malaria parasite transmission

**DOI:** 10.1101/2022.10.18.512810

**Authors:** Jan Stephan Wichers-Misterek, Annika M. Binder, Paolo Mesén-Ramírez, Lilian Patrick Dorner, Soraya Safavi, Gwendolin Fuchs, Tobias L. Lenz, Anna Bachmann, Danny Wilson, Friedrich Frischknecht, Tim-Wolf Gilberger

**Author notes:** contributed equally (Jan Stephan Wichers-Misterek, Annika M. Binder, Paolo Mesén-Ramírez).

## Abstract

Mature gametocytes of *Plasmodium* (*P*.) *falciparum* display a banana (falciform) shape conferred by a complex array of subpellicular microtubules (SPMT) associated to the inner membrane complex (IMC). Microtubule associated proteins (MAPs) define MT populations and modulate interaction to pellicular components. Several MAPs have been identified in *Toxoplasma gondii* and homologues can be found in the genome of *Plasmodium* species, but the function of these proteins for asexual and sexual development of malaria parasites is still unknown. Here we identified a novel subpellicular MAP, termed SPM3, that is conserved within the genus *Plasmodium*., especially within the *Laverania* subgenus, but absent in other Apicomplexa. Conditional knockdown and targeted gene disruption of *Pfspm3* in *P. falciparum* cause severe morphological defects during gametocytogenesis leading to round, non-falciform gametocytes with an aberrant SPMT pattern. In contrast, *Pbspm3* knockout in *P. berghei*, a species with round gametocytes, caused no defect in gametocytogenesis, but sporozoites displayed an aberrant motility and a dramatic defect in sporozoite invasion of salivary glands leading to a decreased efficiency in transmission. Electron microscopy revealed a dissociation of the SPMT from the IMC in *Pbspm3* knockout parasites suggesting a function of SPM3 in anchoring MTs to the IMC. Overall, our results highlight SPM3 as a pellicular component with essential functions for malaria parasite transmission.

**IMPORTANCE:** A key structural feature driving the transition between different life cycle stages of the malaria parasite is the unique three membrane “pellicle”, consisting of the parasite plasma membrane (PPM) and a double membrane structure underlying the PPM termed the “inner membrane complex” (IMC). Additionally, there are numerous linearly arranged intramembranous particles (IMPs) linked to the IMC, which likely link the IMC to the subpellicular microtubule cytoskeleton. Here we identify, localize and characterize a novel subpellicular microtubule associated protein unique to the genus *Plasmodium* (*P*.). The knockout of this protein in the human infecting *P. falciparum* species result in malformed gametocytes and aberrant microtubules. We confirmed the microtubule association in the *P. berghei* rodent malaria homologue and show that its knockout results in a perturbated microtubule architecture, aberrant sporozoite motility and decreased transmission efficiency.

## INTRODUCTION

Despite significant progress in combating malaria, this disease remains a huge burden on health systems in tropical countries worldwide, with an estimated 241 million cases and 627 000 deaths in 2020 (1). The deadliest species, *Plasmodium (P.) falciparum*, accounts for almost 99% of the malaria associated fatalities. *P. falciparum* emerged from the *Laverania* clade, a group of ape infecting *Plasmodium* parasites comprising at least seven cryptic species infecting chimpanzees, gorillas and bonobos (2–5). This clade represents a subgroup within hundreds of known *Plasmodium* species infecting various vertebrate hosts including birds, reptiles, rodents, and humans. Malaria parasites evolved and developed a complex life cycle alternating between invertebrate vectors and vertebrate hosts, leading to the evolution of several highly specialized and morphologically distinct developmental stages that are adapted to a specific niche within different host tissue (6). The drastic morphological changes across the life cycle and the morphogenesis of the different stages are driven by reorganization of cytoskeletal structures (7). A key structural feature driving the transition between different life cycle stages is the unique three membrane “pellicle” of these cells which consists of i) the parasite plasma membrane (PPM) and ii) a double membrane structure, specific to alveolates, termed the “inner membrane complex“ (IMC) or alveoli, which are intramembranous vesicles underlying the PPM and are likely linked to the subpellicular microtubule (SPMT) cytoskeleton (8–11). The different arrangements of the pellicle and its components, especially the IMC and the SPMTs, in structurally distinct malaria parasite stages may have evolved to fulfill the unique requirements for invasion and survival in different host cells (reviewed in (12)). In invasive stages, SPMTs originate at the apical polar ring, an apicomplexan-specific microtubule organizing center (MTOC) and extend to the parasite posterior end in close association with the cytosolic face of the parasite pellicle (11).

The number of SPMTs varies across life cycle stages. For instance, merozoites of *P. falciparum* possess 2–4 SPMTs (14, 15), whereas ookinetes have 60, sporozoites 14 and gametocytes 21 SPMTs (11). SPMTs of *P. falciparum* are extremely stable exemplified by their resistance to classic microtubule depolymerizing agents (16). Recent studies showed polyglutamylation of SPMTs in merozoites (17) and gametocytes (13) and suggested its involvement in SPMT stability (17). A second population of microtubules, the spindle microtubules, is necessary to coordinate chromosome segregation and these extend from a single MTOC throughout the nucleus (14, 18). Gametocytes, the blood-circulating sexual stages critical for malaria transmission, also possess an elaborate IMC and an array of SPMTs (19–22). Strikingly, mature gametocytes of *P. falciparum* and the other species of the clade *Laverania* display a banana (falciform) shape while round ameboid gametocytes are observed for many rodent-infecting species such as *P. berghei* or in other human-infecting species like *P. vivax* (22–25). The underlying molecular differences between species that define this morphology are not clearly understood. The elongated cell shape of *P. falciparum* gametocytes is conferred by the assembly of an array of SPMTs recruited to the nascent IMC in early gametocyte stages which extend laterally as gametocytes mature (20). Here, the SPMTs are 27 to 38 nm in diameter (26) and linearly spaced 10-25 nm apart (22, 24). In contrast to invasive stages such as sporozoites, merozoites and ookinetes, gametocytes lack apical polarity and an apical polar ring from which SPMTs emanate. A recent study reports the nucleation of SPMTs in gametocytes at the outer centriolar plaque, a non-mitotic MTOC embedded in the nuclear membrane of the parasite (13). The SPMT network disassembles in stage V gametocytes, providing an increased deformability that is expected to permit transmigration across the endothelial barrier and exit from the bone marrow (27, 28). Although robust gametocyte motility *in vivo* has not been reported, their elongated morphology, which is similar to motile “zoite” forms, may contribute to movement between tissues and vessels (29).

Numerous studies in the model apicomplexan organism *Toxoplasma (T.) gondii*,identified several microtubule-associated proteins (MAPs) decorating the SPMTs including *Tg*SPM1/2 (30), *Tg*TrxL1/2, and *Tg*TLAP1/2/3/4 (31, 32). MAPs are expected to define SPMT subpopulations with different properties and to modulate microtubule stability, mechanical properties and resistance to depolymerization (30, 32–35). Some of these MAPs, such as *Tg*TrxL1, *Tg*TrxL2 and *Tg*SPM1, are known to localize within the lumen of SPMTs (36) while others might interact with their external surface. Amino acid repeats in the *Tg*MAPs are important for microtubule localization and tubulin binding (30). *Tg*SPM1 and *Tg*TrxL1 are important for stability of the SPMTs, but are not essential for parasite development, suggesting remarkable functional redundancy (30, 36). *Tg*AC9 and *Tg*AC10 were shown to be required for the organization of the SPMTs at the apical cap and mediate interaction to IMC components (37), while *TgDCX* has been shown to modify the structure of tubulin in the conoid (38).

Homologues for most of the *Tg*MAPs can be found in the genome of *Plasmodium* species, but none has been experimentally characterized for either asexual or sexual developmental stages so far. The presence of *Plasmodium* MAPs within the SPMT lumen has been suggested previously (39, 40), but SPMT structures resembling those expected for *Pf*SPM1, *PfSPM2* and *Pf*TRXL1 in SPMT lumen have only been identified recently by cryoEM and modelling, supporting that these proteins have a role in SPMT function (26).

Although the findings of Ferreira *et al*. (26) and the presence of several homologues to *T. gondii* MAPs in the *P. falciparum* genome suggest that potentially many *Pf*MAPs might have similar functions to those in *T. gondii*, differences in the malaria parasite life cycle and the structure of microtubules open up the possibility that they have evolved MAPs of unique function.

Here we identify SPM3 (PF3D7_1327300 / PBANKA_1342500) as a novel MAP that is highly conserved within *Plasmodium*, especially within the *Laverania* clade, but absent in other Apicomplexa such as *T. gondii*. Characterization of SPM3 reveals its importance in falciform gametocyte morphology and sexual development in *P. falciparum*, as well as its role in the transmission of *P. berghei* sporozoites.

## RESULTS

### *Pf*SPM3 is a *Plasmodium-specific* microtubule-associated protein in *P. falciparum* merozoites

PF3D7_1327300 was initially identified as a putative interaction partner of the IMC protein *Pf*PhIL1 (41). Phylogenetic analysis revealed orthologous proteins of *Pf*SPM3 with high similarity (>89% amino acid similarity) in *P. reichenowi* and *P. gaboni* (Figure 1A). Other *Plasmodium* species also appear to have homologous proteins, but with significantly lower similarity (<50% amino acid similarity) than in *Laverania* parasites. The only homologous protein sequence outside of the genus *Plasmodium* was found in *Hepatocystis* spp. (Figure 1A).

**Figure 1:**
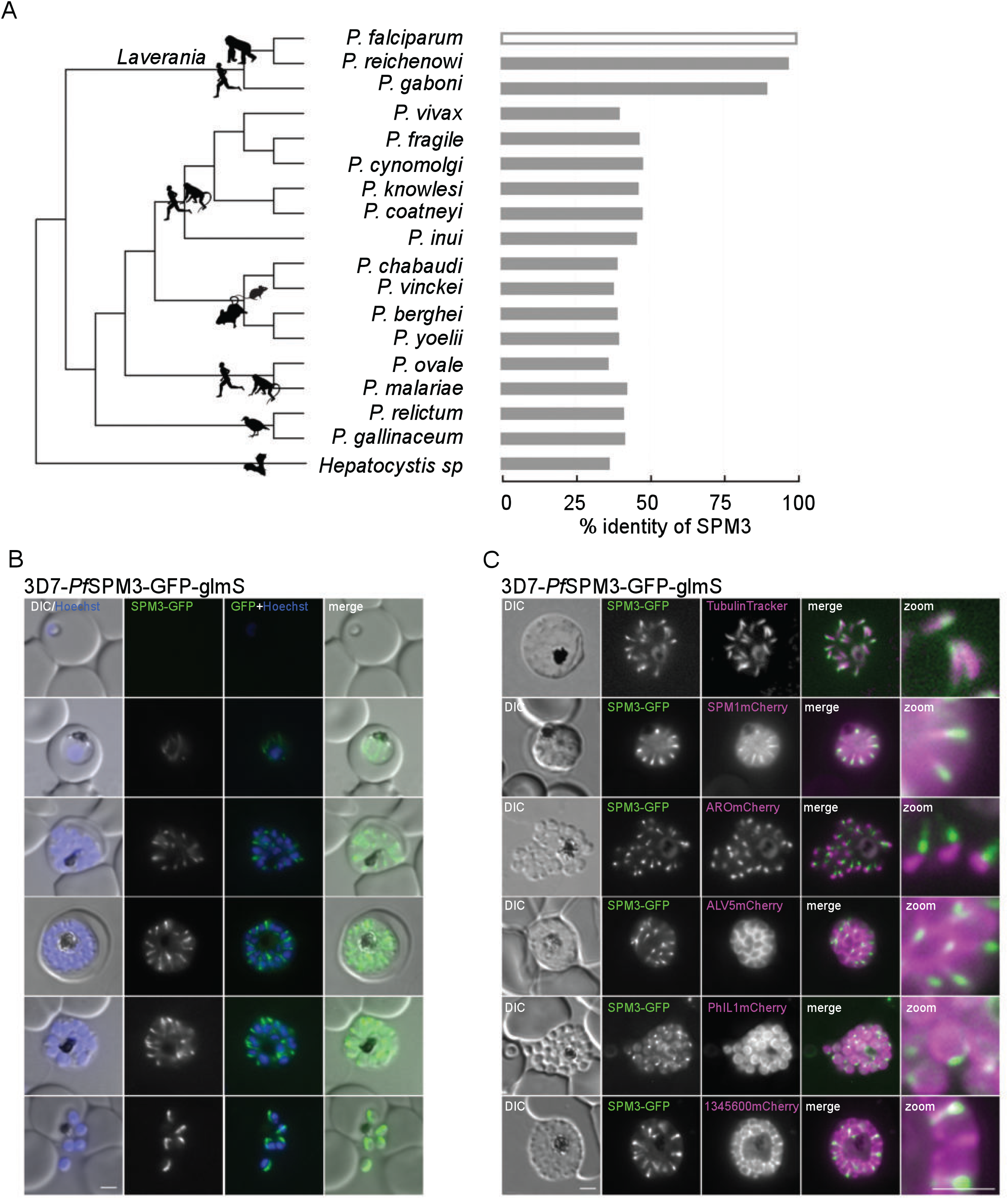
*Pf*SPM3 localizes to the SPMTs of *P. falciparum* merozoites. **(A)** Phylogenetic relatedness of sequences homologous to SPM3 across species (fast minimum evolution tree based on Grishin protein distance), and similarity along homologous sequence stretches. Silhouettes (from phylopic.org) depict representatives of the vertebrate hosts for each lineage. **(B)** Localization of *Pf*SPM3-GFP by live-cell microscopy during intracellular development cycle in 3D7-*Pf*SPM3-GFP-glmS. **(C)** Co-localization of *Pf*SPM3-GFP with TubulinTracker, microtubule associated protein *Pf*SPM1mCherry, the rhoptry marker protein ARO-mCherry, and the IMC marker proteins ALV5mCherry, PhIL1mCherry and PF3D7_ 1345600mCherry. Nuclei were stained with Hoechst-33342. Scale bar, 2 μm. Zoom factor 400% with scale bar of 1 μm.

To test putative IMC association and to investigate the physiological role of this protein in *P. falciparum*, we first generated transgenic parasites expressing a C-terminal GFP tag. Genomic integration of a SLI-based plasmid (42) into the *spm3* locus was verified by PCR (Figure S1A). Expression and localization of the fusion protein was analyzed by fluorescence microscopy. The GFP-fusion protein was detectable from trophozoites stage onwards in agreement with its transcriptional profile (43, 44). Unexpectedly, its localization did not resemble the localization of any IMC resident proteins but was reminiscent of SPMT distribution (45) (Figure 1B). Localization of GFP-tagged *Pf*SPM3 at the microtubules was visualized by co-localization with TubulinTracker (Figure 1C, first row), showing partial overlap with a more pronounced signal of *Pf*SPM3 towards one end of the SPMT. This SPMT association was substantiated by co-localization with an overexpressed mCherry-tagged *Pf*SPM1 (PF3D7_0909500) (Figure 1C, second row), the homologue of *Tg*SPM1 (30), which showed a similar distribution to the TubulinTracker with some cytosolic background.

Co-localization of *Pf*SPM3-GFP with *Pf*SPM1-mCherry or TubulinTracker, respectively, underlined the more polar distribution of *Pf*SPM3 along SPMTs compared with *Pf*SPM1-mCherry and TubulinTracker. Based on this localization data, PF3D7_1327300 was named *Pf*SPM3 – subpellicular microtubule protein 3 – acknowledging the previously published SPMT associated proteins *Tg*SPM1 and *Tg*SPM2 in *T. gondii* (30). To confirm the polar distribution of *Pf*SPM3-GFP on the SPMT, we co-localized the protein with the rhoptry bulb marker ARO-mCherry (46, 47) and found that the *Pf*SPM3-GFP signal is significantly concentrated towards the apical pole of the SPMT in nascent merozoites (Figure 1C, third row). Finally, in order to investigate the previously described potential interaction of *Pf*SPM3 with *Pf*PhIL1 (41), *Pf*SPM3 was co-localized with three different IMC marker proteins: ALV5-mCherry (48, 49), PhIL1-mCherry (20, 50, 51) and PF3D7_1345600 mCherry (52) (Figure 1B, rows 4-6). Co-localization with these IMC markers shows, in agreement with the *Pf*SPM1 and TubulinTracker investigations, that *Pf*SPM3 lies in close proximity, but spatially distinctive, from the IMC.

### *Pf*SPM3 is dispensable for intraerythrocytic development

To assess the function of *Pf*SPM3 in asexual development of *P. falciparum* parasites, a glucosamine-inducible degradation system was used to degrade *Pf*SPM3 mRNA (glmS ribozyme system (53)) (Figure S1A, B). The glmS ribozyme was introduced upstream of the 3’-untranslated region in the modified *spm3* locus. Although conditional knockdown of *Pf*SPM3-GFP upon addition of 2.5 mM glucosamine resulted in reduction of the GFP signal (Figure 2A) it had neither measurable effect on parasite proliferation during asexual blood stage parasite development (IDC) (Figure 2B) nor on schizont morphology (Figure 2A). To exclude that residual amounts of *Pf*SPM3 upon knockdown might be sufficient for parasite development we tested an additional approach using targeted gene disruption (TGD) (42) (Figure S1B). Successful integration and consequently disruption of the *Pfspm3* gene was validated by PCR (Figure S1A). Expression and localization of the truncated protein fragment was monitored by fluorescence microscopy (Figure 2C). The engineered deletion (aa 137–1796) of *Pf*SPM3 resulted in a cytosolic GFP signal of the truncated *Pf*SPM3-TGD fragment, but as before, these *Pf*SPM3 deficient parasites showed neither a growth nor a morphological phenotype (Figure 2C, D). Congruently, these parasites revealed an indistinguishable SPMT and IMC architecture compared to wild-type parasites as visualized by co-expression of the IMC marker proteins ALV5-mCherry (48, 49) and PF3D7_1345600-mCherry (52) or by co-staining with TubulinTracker (Figure 2E) confirming the dispensability of *Pf*SPM3 for asexual blood stage development.

**Figure 2:**
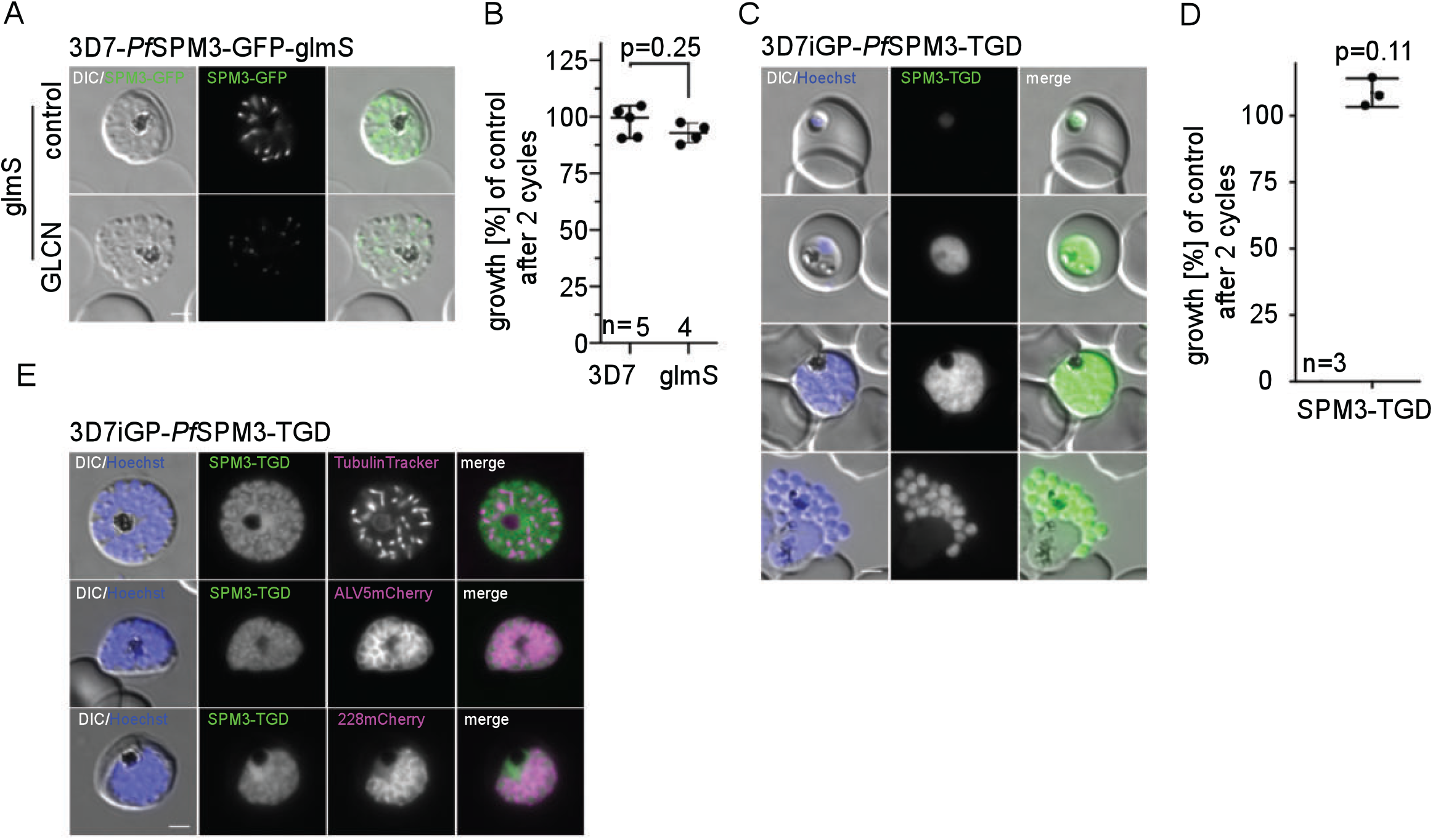
*Pf*SPM3 is dispensable for the asexual replication cycle. **(A)** Representative live-cell microscopy of 3D7-*Pf*SPM3-GFP-glmS schizonts cultured either with or without (control) 2.5 mM glucosamine (GLCN) at 40h post addition of GLCN. Scale bar, 2 μm**. (B)** Growth of 3D7 and 3D7-*Pf*SPM3-GFP-glmS parasites treated with or without 2.5 mM GLCN after two parasite replication cycles as determined by flow cytometry. Shown are relative parasitemia values, which were obtained by dividing the parasitemia of GLCN-treated cultures by the parasitemia of the corresponding untreated ones. Displayed are means +/− standard deviation (SD) of independent growth experiments with the number of experiments (n) indicated. P-values displayed were determined with a two-tailed unpaired t-test with Welch’s correction. **(C)** Localization of truncated *Pf*SPM3-TGD-GFP fusion protein by live-cell microscopy across the IDC in the 3D7-iGP background. **(D)** Growth curves of 3D7-iGP-*Pf*SPM3-TGD vs. 3D7-iGP parasites after two parasite replication cycles as determined by flow cytometry. Three independent growth experiments were performed, and a summary is shown as percentage of growth compared to the parental parasite line. Displayed are means +/− standard deviation (SD) of independent growth experiments with the number of experiments (n) indicated. P-values displayed were determined with a one-sample t test. **(E)** Co-localization of the truncated *Pf*SPM3-TGD-GFP fusion protein with TubulinTracker and the IMC marker proteins ALV5mCherry and PF3D7_ 1345600mCherry. Nuclei were stained with Hoechst-33342. Scale bar, 2 μm. Zoom factor 400% with scale bar of 1 μm.

### *Pf*SPM3 is important for falciform gametocyte morphology

In addition to the 3D7-based transgenic parasites, we generated a corresponding cell line on the 3D7-iGP (54) background. This cell line (3D7-iGP-*Pf*SPM3-GFP-glmS) allows the robust induction of sexual commitment of the parasite and therefore enables the localization and assessment of *Pf*SPM3 function during gametocytogenesis. As expected, fluorescence microscopy revealed close SPMT localization of *Pf*SPM3 in gametocytes (Figure 3A). Of note, the *Pf*SPM3-GFP signal disappears earlier than the TubulinTracker signal, indicating loss of detectable *Pf*SPM3 prior to the disassembly of the SPMTs in late gametocyte stages (IV/V gametocytes, Figure 3A). Conditional knockdown via the glmS system resulted in significantly reduced numbers of gametocytes (mean of 52.6% ± 16.1% at day 10 pi) compared to control cultures (Figure 3B). Gametocytes were morphologically distinct and appeared as malformed (round) and non-falciform gametocytes. Importantly, these *Pf*SPM3 depleted gametocytes revealed an aberrant SPMT pattern (Figure 3A, C). To exclude detrimental effects of glucosamine on gametocyte development and SPMT architecture, we assessed the phenotype in the control 3D7-iGP-*Pf*SPM3-GFP parasites with an inactive M9 mutant ribozyme sequence (53). No difference in gametocyte morphology or gametocytemia was observed in these parasites in the presence of 2.5 mM glucosamine (Figure 3B, C and S2A). The phenotype was further confirmed by using the 3D7-iGP-*Pf*SPM3-TGD parasite cell line that expresses the truncated, cytosolic *Pf*SPM3. As for the glmS-based approach, after induction of gametocytogenesis the progression and maturation of gametocytes was heavily impaired resulting in the absence of falciform gametocytes and leading to the formation of malformed, round gametocytes with aberrant SPMT organization. (Figure 3D, E and S2B). Altogether these results indicate an essential function of *Pf*SPM3 in gametocyte development and the falciform morphology of *P. falciparum* gametocytes.

**Figure 3:**
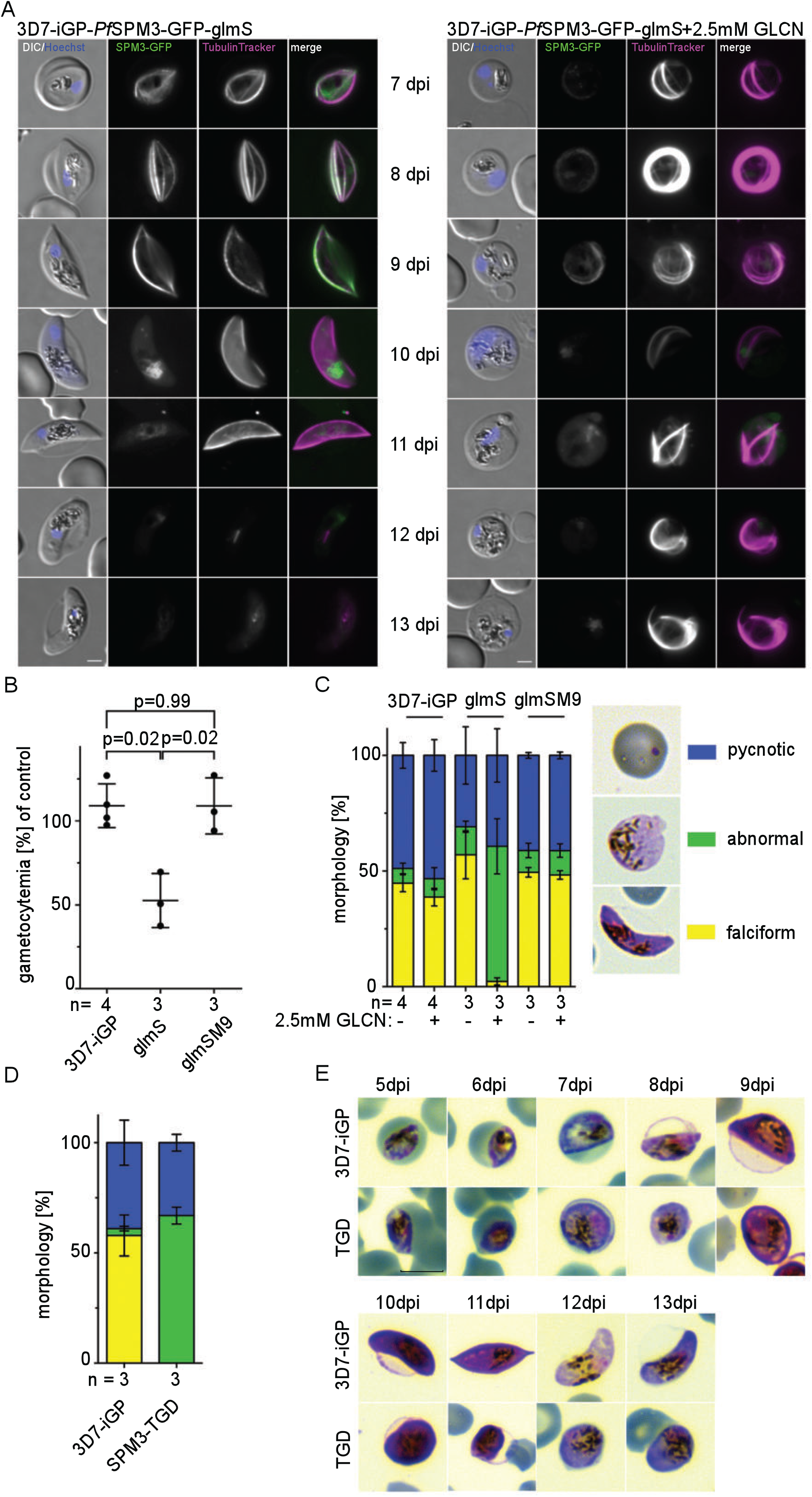
*Pf*SPM3 deficiency interferes with gametocytogenesis and leads to non-falciform morphology. **(A)** Live-cell microscopy of 3D7-iGP-*Pf*SPM3-GFP gametocytes (day 7 – day 13 post gametocyte induction) cultured either with or without (control) 2.5 mM glucosamine (GLCN). Microtubules were visualized with TubulinTracker and nuclei were stained with Hoechst-33342. Scale bar, 2 μm. **(B)** Gametocytemia at day 10 post gametocyte induction was determined by counting between 1,096–2,632 (mean 1,793) cells per condition in Giemsa-stained thin blood smears. The relative gametocytemia values (%) displayed were obtained by dividing the gametocytemia of GLCN-treated cultures by the gametocytemia of the corresponding untreated cultures. Displayed are means +/− SD of independent growth experiments with the number of experiments (n) indicated. A two-tailed unpaired t-test with Welch’s and Benjamini-Hochberg correction was used to calculate multiplicity adjusted p-values for 3D7-iGP-*Pf*SPM3-GFP-glmS or 3D7-iGP-*Pf*SPM3-GFP-glmSM9 versus 3D7-iGP parasites all cultured with 2.5 mM GLCN. **(C)** Classification and quantification of gametocyte morphology within parasite populations of 3D7-iGP, 3D7-iGP-*Pf*SPM3-GFP-glmS and 3D7-iGP-*Pf*SPM3-GFP-glmSM9 cultured either with or without (control) 2.5 mM GLCN. Between 589–2632 erythrocytes and 19–69 (mean 48) parasites per condition in Giemsa-stained thin blood smears were analyzed. Yellow = falciform, blue = pycnotic and green = abnormal; representative images are shown in the right panel) at day 10 post gametocyte induction. **(D)** Quantification of gametocyte morphology (yellow = falciform, blue = pycnotic and green = abnormal) at day 10 post gametocyte induction for 3D7-iGP (control) and 3D7-iGP-SPM3-TGD parasites. For each condition, the proportion of parasite stages in erythrocytes was determined in three independent experiments (total number of cells screened: n_3D7-iGP_= 2,694/1,431/2,742 and n_3D7-iGP-SPM3-TGD_ = 1,328/1,144/925) and displayed as a percentage. **(E)** Representative Giemsa smears of 3D7-iGP and 3D7-iGP-*Pf*SPM3-TGD stage II–V gametocytes (day 5 – day 13 post gametocyte induction). Scale bar 5 μm.

### SPM3 deletion in *P. berghei* impacts sporozoite motility and transmission by mosquitoes

To extend our functional investigation into the role of SPM3 in mosquito stages, we turned to the rodent model parasite *P. berghei*, for which the full life cycle is readily available. We first generated a *P. berghei* parasite line where the orthologous gene of *P. falciparum spm3* (*Pb*SPM3, PbANKA_1342500) was tagged with GFP (*Pb*SPM3-GFP) (Figure S3A, B). *Pb*SPM3-GFP was detectable in oocysts, as well as in midgut and salivary gland sporozoites. In sporozoites, the fluorescent signal extended from the apical end towards the rear around the nucleus (Figure 4A and S4A). Subsequent co-staining with an anti-tubulin antibody revealed co-localization of *Pb*SPM3-GFP with tubulin (Figure 4A) in midgut and salivary gland sporozoites. C-terminal tagging of *Pb*SPM3 with GFP did not affect parasite life cycle progression, as SPM3-GFP parasites showed normal midgut infection and midgut oocyst numbers as well as numbers of both midgut and salivary gland sporozoites comparable to wild-type (Figure S4B, C)

**Figure 4:**
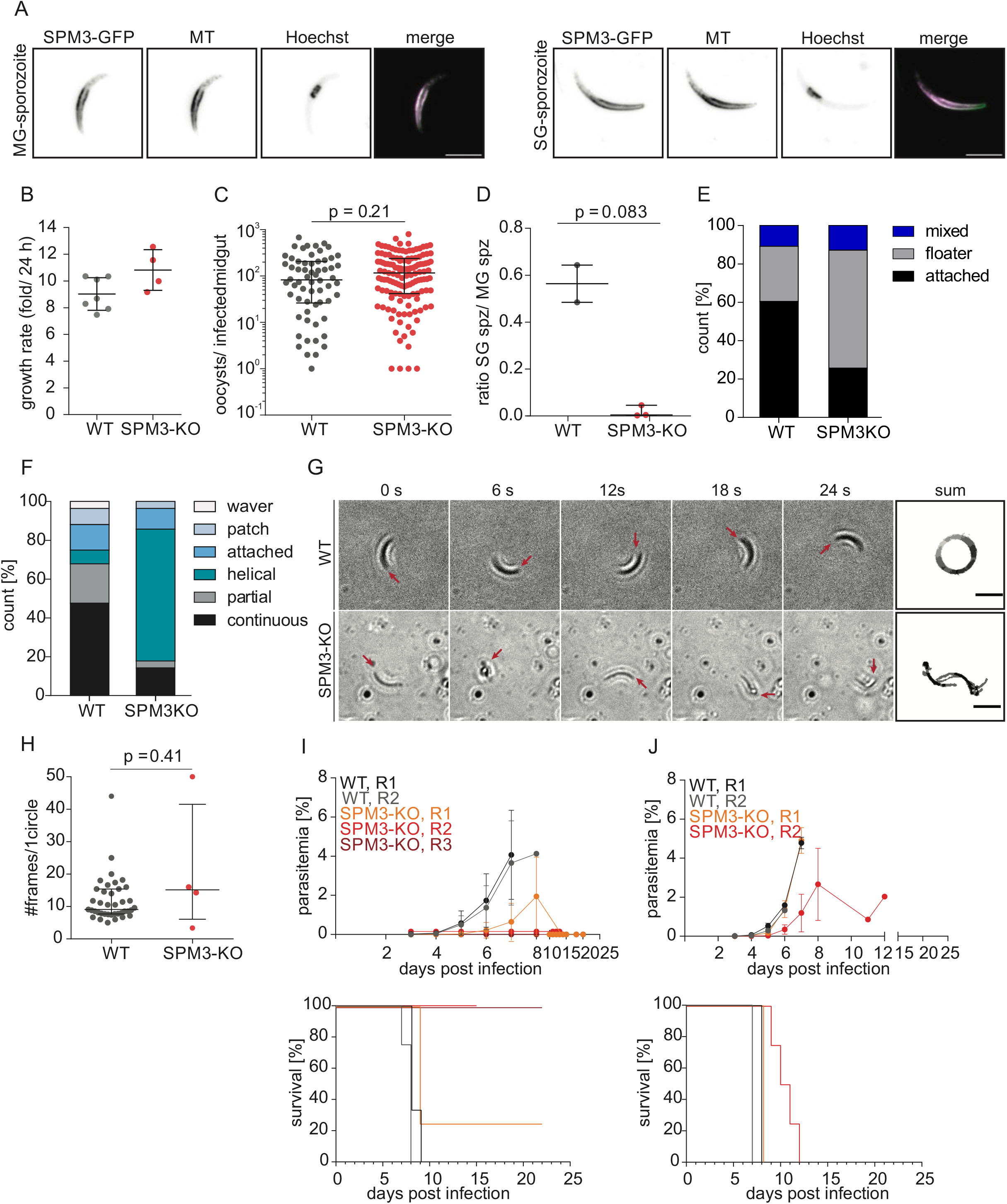
*Pb*SPM3 localizes to SPMTs and is critical for sporozoite motility, salivary gland invasion and transmission to mammalian host. **(A)** *Pb*SPM3 localizes along microtubules in midgut and salivary gland sporozoites. Representative images of fixed sporozoites. Shown are 2D projections of an acquired Z-stack. Microtubules (MT) were stained with an anti-tubulin antibody. Nuclei were stained with Hoechst-33342. Merge image with microtubules in magenta and *Pb*SPM3-GFP in green. Scale bar, 5 μm. **(B)** Deletion of *pbspm3* does not affect asexual blood-stage growth rate. Asexual blood stage growth rates were calculated based on the parasitemia value at day 8 post-infection with a single infected red blood cell (iRBC) intravenously. PbANKA wild-type (WT) growth rates was plotted as reference and has been determined and previously published by our laboratory (94). Mean fold change within a single parasite replication cycle (24 hours for *P. berghei*) with SD is indicated. **(C)** Similar oocyst numbers and infection rates of wild-type and *Pb*SPM3-KO-infected *A. stephensi*. Shown are pooled data from two (WT) and three (*Pb*SPM3-KO) independent cage feeds. A total of 79 (WT) and 151 (*Pb*SPM3-KO) mosquitoes were analyzed on day 11 and day 12 post infection as technical replicates of which 78.5% (62) and 88.1% (133) were infected, respectively (p = 0.08, Fisher’s exact test). For statistical analysis comparing oocyst numbers in infected midguts, Mann-Whitney test was performed. Black line indicates median with error bars representing interquartile range (IQR). **(D)** Ratio of salivary gland resident versus midgut resident sporozoites determined at day 18 and 20 post infection from two (WT) and three (*Pb*SPM3-KO) independent cage infections. P-value determined with unpaired t-test with Welch’s correction. Black line indicates median with error bars representing interquartile range. **(E)** Percentage of wild-type and *Pb*SPM3-KO sporozoites isolated from salivary glands displaying movement after attachment or floating in the medium. Pooled data from two (WT) with 139 sporozoites and three (*Pb*SPM3-KO) with 109 sporozoites independent cage feeds with two technical replicates per cage infection. For statistical analysis comparing differences in gliding categories, a Chi-square test was performed. **(F)** Different types of movement patterns observed in sporozoites analysed in E. **(G)** Wild-type like circular (left panel) and helical (right panel) movement pattern. Numbers indicate time in seconds. Red arrows point to the apical end of the sporozoite. Scale bar: 10 μm. **(H)** Sporozoite speeds from continuously gliding wild-type and *Pb*SPM3-KO sporozoites as shown in F. As the *Pb*SPM3-KO mutant had only four continuously gliding sporozoites in the total sporozoites analyzed from the three biological replicates, only four data points are shown in comparison to n = 40 for wild-type. P-value determined with Mann-Whitney test. Black line indicates median with IQR. **(I)** Parasitemia curves and Kaplan-Meier plots showing increase in blood stage parasitemia (left) and mouse survival after natural transmission. 2 and 3 independent infections for wild-type and *Pb*SPM3-KO, respectively, using 3–4 mice per experiment (in total: WT n=7, *Pb*SPM3-KO n=12). Wild-type is shown in grey and black; *Pb*SPM3-KO in different shades of red; see also Table 1. Note that mice from two replicates remained negative over the course of parasitemia monitoring and hence parasitemia curve of replicate 3 was slightly nudged to allow for better visibility. Also, lines in Kaplan-Meier plots in J and I were slightly nudged to allow for better visibility. **(J)** Parasitemia curves and Kaplan-Meier plots showing increase in blood stage parasitemia (left) and mouse survival after injection of 1,000 sporozoites intravenously. Two independent infections for wild-type (grey, black) and *Pb*SPM3-KO (red, orange) using 3–4 mice per experiment (in total: WT n=7, *Pb*SPM3-KO n=8).

In order to probe into *Pb*SPM3 function, we generated a clonal *Pb*SPM3 knockout (*Pb*SPM3-KO) cell line (Figure S3C, D). Similar to the *P. falciparum* gene disruption, no apparent difference in blood stage growth was detected for *Pb*SPM3-KO compared to our previously determined growth rates of *P. berghei* wild-type parasites (55, 56) (Figure 4B). Next, to investigate the progression of this parasite line in mosquitoes we fed naïve *Anopheles stephensi* mosquitoes on mice infected with *Pb*SPM3-KO parasites. Surprisingly, we found that infection and colonization of the mosquito midgut by *Pb*SPM3-KO parasites was similar to wild-type parasites, as measured by the percentage of fed mosquitoes that had oocysts and the median numbers of oocysts per infected mosquito (*Pb*SPM3-KO: 116 (IQR: 42–238); WT: 83 (IQR: 26–206) (Figure 4C). Furthermore, we found similar numbers of *Pb*SPM3-KO sporozoites (median: 83,000, IQR: 49,000–94,000) compared to wild type (median: 39,000, IQR: 30,000–63,000) maturing in mid-gut oocysts (Table1). These findings indicate that neither gametocyte development, exflagellation, ookinete development, nor midgut traversal are significantly affected by *Pb*SPM3 gene knockout.

**Table 1:** Numbers and infectivity of *Pb*SPM3-KO sporozoites compared to wild-type (WT)

| Parasite line | Midgut spz / mosquito<br>±SD (replicates) | Salivary gland spz / mosquito<br>±SD (replicates) | 1,000 spz i.v. |  | natural (by bite) |  |
| --- | --- | --- | --- | --- | --- | --- |
|  |  |  | Prepatency time<br>(#infected / #total) | Parasitemia at d6<br>(infected only) | Prepatency time<br>(#infected / #total) | Parasitemia at d6<br>(infected only) |
| WT | 34,000 ± 6,000<br>55,000 ± 16,000 | 21,000 ± 1,000<br>27,000 ± 10,000 | 3.6 (7/7) | 1.4 % | 3.7 (7/7) | 1.5 % |
| *PbSPM3*-KO | 70,000 ± 22,000<br>90,000 ± 9,000<br>67,000 ± 18,000 | 3,000 ± 1,000<br>200 ± 40<br>300 ± 50 | 4.6 (8/8) | 0.8 % | 5.7 (3/12) | 0.3 % |

n = 2 for WT, n = 3 for *PbSPM3*-KO for by bite transmission and n = 2 for *PbSPM3*-KO for i.v. injection Parasitemia values given for all mice that became positive within 20 days; C57/BL6 mice were used.

Next, we tested whether *Pb*SPM3-KO might affect the further development of sporozoites. While salivary glands of mosquitoes infected with wild-type parasites showed over 20,000 sporozoites (median: 21,000, IQR: 18,000–34,000), only a few hundred were found for *Pb*SPM3-KO sporozoites (median: 300, IQR: 220–2,690) indicating a drastic (99%) decrease in salivary gland over midgut sporozoite ratio (Figure 4D, Table 1).

To pinpoint the cause for the reduced number of *Pb*SPM3-KO sporozoites in salivary glands we assessed the effect of *Pb*SPM3 deletion on sporozoite motility, an essential asset for the parasite entry into salivary glands and for establishing infection in human liver cells. Wild-type *P. berghei* sporozoites isolated from salivary glands attached and migrate on flat surfaces in a counter-clockwise circular manner at speeds exceeding 1 micrometer per second (40). While 60%of the wild-type sporozoites attached and moved on the glass coverslip most of *Pb*SPM3-KO sporozoites (61%) remained floating in the medium (Figure 4E). For the majority of *Pb*SPM3-KO sporozoites that managed to attach and move, a striking new phenotype of sporozoite locomotion was revealed: the parasites moved in a helical way (Figure 4F, G) and deviated from the circular path observed in most of the wild-type sporozoites. The few *Pb*SPM3-KO sporozoites that showed persistent circular gliding motility moved with a similar speed to wild-type sporozoites (Figure 4H).

To further dissect the physiological consequences of this aberrant cell motility, we investigated the transmission capacity of mosquitoes infected with *Pb*SPM3-KO parasites. We found that only 4 out of 12 mice became infected after being bitten by *Pb*SPM3-KO infected mosquitoes, while all mice bitten by mosquitoes infected with wild-type parasites became infected (Figure 4I, Table 1). Furthermore, we noticed a delay in appearance (prepatency) of 2 days in mice that did became infected with *Pb*SPM3-KO parasites. This drop in infectivity is likely due to the much lower load of sporozoites within the salivary glands. We next tested mouse liver infectivity of *Pb*SPM3-KO sporozoites by direct intravenous inoculation using direct inoculation of an equal number of purified sporozoites, because the lower level of sporozoites in salivary glands made direct transmission from mosquito to mice unreliable for this assay (57). All mice infected through the intravenous injection of 1,000 *Pb*SPM3-KO sporozoites became infected (Figure 4J). These results suggest that the reduced number of *Pb*SPM3-KO sporozoites in the salivary glands and their aberrant motility affects the transmission from the mosquito to the mammalian host, but not the invasion of liver cells.

### *Pb*SPM3 deletion disturbs SPMT architecture in *P. berghei*

To identify the cause for the aberrant motility of *Pb*SPM3-KO sporozoites we investigated the SPMT organization in fixed midgut sporozoites by transmission electron microscopy. Preparation of thin sections from oocysts allows the imaging of sporozoites in cross sections (Figure 5A). We found that near the apical end of *Pb*SPM3-KO sporozoites, the arrangement of SPMTs largely followed that of wild-type sporozoites, with a single SPMT being somewhat apart from the others and all SPMTs closely associated with the inner membrane complex (Figure 5A). However, we found that further away from the apical end (as discernible by an increased sporozoite diameter, the presence of apical organelles such as micronemes and rhoptries and an increased inter-microtubule distance), the SPMTs were not associated with the IMC and often appeared deep within the cytoplasm (Figure 5A, B). This suggests a dissociation from the IMC. We next quantified the distance between SPMTs and IMC by measuring from microtubule center towards the inner side of the IMC. Also, the SPMT-IMC distance was increased in the SPM3-KO with only 1% of SPMTs being more than 40 nm apart from the IMC in wild type compared to 19 % in *Pb*SPM3-KO (Figure 5C, S4D). These observations points towards a role of *Pb*SPM3 in anchoring the SPMT to the IMC in sporozoites.

**Figure 5:**
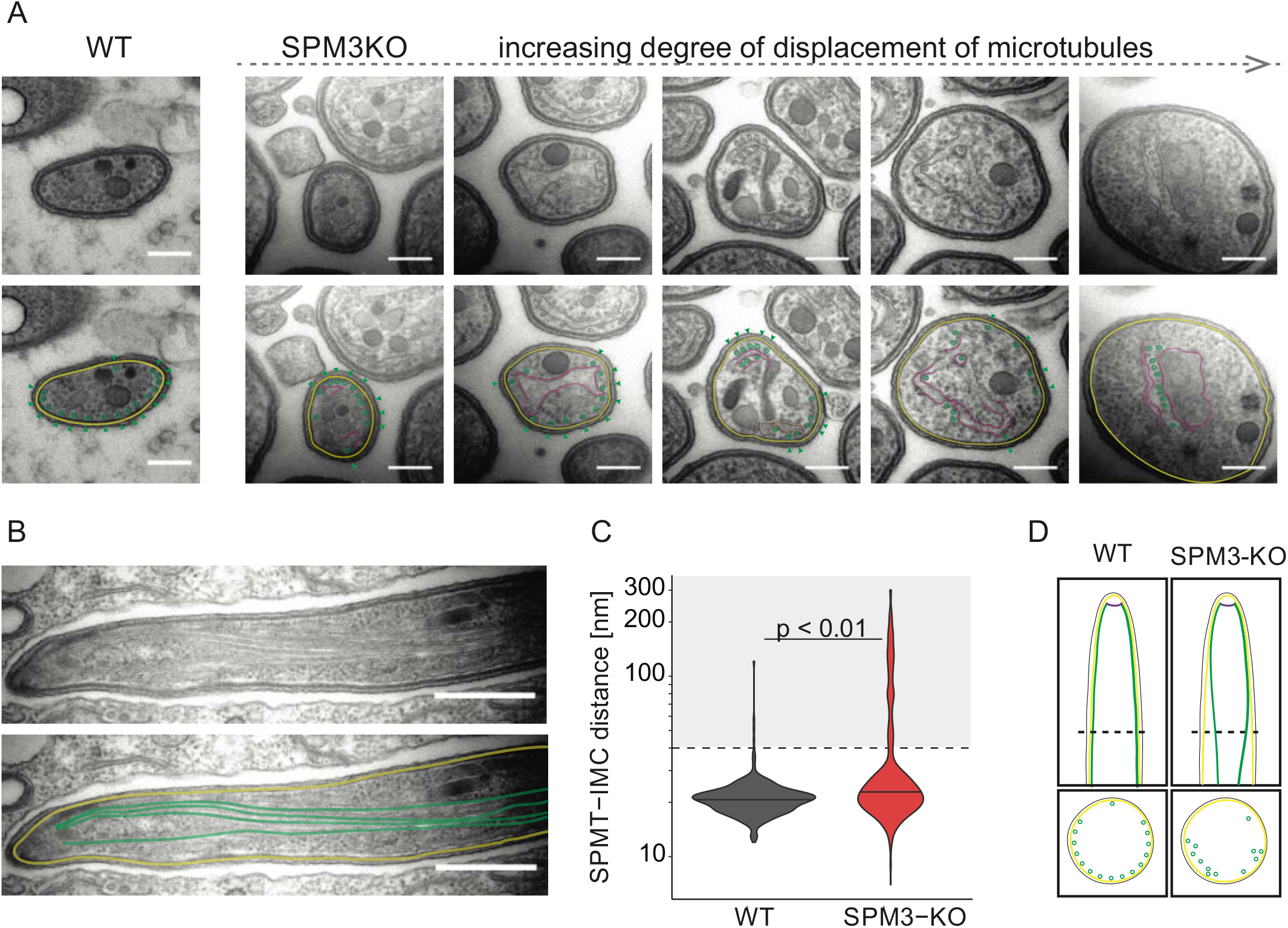
SPMT dissociate from the IMC in *Pb*SPM3-KO sporozoites. **(A)** Transmission electron micrographs showing cross sections through wild-type (WT, left panel) and *Pb*SPM3-KO (right panel) *P. berghei* sporozoites within oocysts at day 15 post mosquito infection. Single images are ordered according to an increased degree of microtubule displacement. Scale bar, 200 nm. Color code: green: SPMTs, yellow: IMC, magenta: membrane of unknown origin. **(B)** Longitudinal section through a *Pb*SPM3-KO midgut sporozoite. Note the dissociation of the microtubules from the IMC with increasing distance from the apical end. **(C)** Distance between microtubules and IMC with black line indicating median. The dashed line indicates the preset cut-off value of 40 nm distance as distances equal or below 40 nm were defined as IMC-close microtubules according to the findings in microtubule-IMC distance by Ferreira *et al*. (26). Grey background highlights all values being not considered IMC-close with the corresponding percentages above. In total 349 wild-type and 579 *Pb*SPM3-KO microtubules were measured. Spread of microtubule-IMC distance was statistically analyzed using a linear mixed model. **(D)** Proposed model showing the dissociation of the microtubules from the IMC with increasing distance from the apical end in the *Pb*SPM3-KO. Dashed-line in the cartoon in the longitudinal section shows the position of the cross-section shown below.

## DISCUSSION

In this study we identified SPM3, a novel *Plasmodium* specific microtubule-associated protein critical for gametocyte development and morphology of *P. falciparum* gametocytes and motility of *P. berghei* sporozoites. *Pf*SPM3 is a high molecular weight protein with no predicted transmembrane domain initially identified as a potential IMC component by *Pf*PhIL1-based proximity dependent biotinylation in schizonts (41) and by *Pf*BLEB-based proximity-dependent biotinylation in stage II–III gametocytes, a protein associated with the basal complex of the parasite (58). By using co-localization approaches we showed that *Pf*SPM3 co-localize with the SPMTs in asexual and sexual stages and only partially with the IMC compartment. In asexual stages, *Pf*SPM3 appears to be more pronounced at one pole of the SPMTs compared to *Pf*SPM1, a homologue of a well-characterized *T. gondii* MAP, which showed a homogenous pattern along the microtubule. MAPs display restricted localization patterns along the SPMTs and may compartmentalize different regions of the microtubule by interacting with pellicular structures such as the IMC (32). Of note, similar to the pronounced apical localization observed for *Pf*SPM3, *Tg*TLAP3 has been shown to selectively coat the SPMTs in the apical cap region and the intraconoidal microtubules of both mother and daughter cells (32).

*Pf*SPM3 appears to be – in agreement with genome-wide mutagenesis screens (59, 60) – dispensable for asexual blood stage proliferation since depletion of the protein showed no growth effect, indicating redundancy of MAPs or a stage-specific function. In accordance with the latter, we provide experimental evidence that SPM3 is a critical factor for gametocyte development and morphology in *P. falciparum*. Depletion of *Pf*SPM3 led to an aberrant organization of the SPMTs in gametocytes, a reduction in the number of viable gametocytes and to the formation of round gametocytes differing from the typical falciform shape of *P. falciparum*.

The drastic morphological transformation observed during gametocytogenesis from a committed ameboid trophozoite to a banana-shaped, falciform gametocyte is orchestrated by the expansion of the IMC network and the extension of SPMT arrays (22). These pellicular components maintain the elongated shape of *P. falciparum* and other *Laverania* gametocytes (24). SPMTs also confer rigidity to the immature gametocytes, which is proposed to be involved in the sequestration of gametocytes in the microvasculature and tissues of the human host to avoid splenic clearance (25). Later depolymerization and disassembly of the SPMT is expected to increase cellular deformability and release into the bloodstream where gametocytes are taken up by the feeding mosquitoes. The falciform morphology is the characteristic hallmark of *Plasmodium* species within the *Laverania* clade and is expected to favor the transmigration from mature gametocytes from the bone marrow reservoir to the bloodstream (22, 24). Nevertheless, the molecular components that determine this morphology remain largely unknown. The phenotype observed in gametocytes upon depletion of *Pf*SPM3 and the low conservation of the protein outside the subgenus *Laverania* could indicate that *Laverania* SPM3 proteins are somehow involved in the falciform architecture of the late-stage gametocytes. Consistent with this notion, we observed no evident phenotype in the roundish *P. berghei* gametocytes from rodents. Previous studies with non-falciform round gametocytes of *P. berghei* and *P. knowlesi* show a discontinuous IMC network and pellicle architecture that does not surround these stages (66). This reflects the pronounced structural differences in gametocyte development between the rodent-infecting *P. berghei* and the human-infecting *P. falciparum* parasites (25). The stage-specific phenotypes observed in *P. berghei* and *P. falciparum* indicates that in the different species MAPs can be repurposed according to requirements of the different stages and might be explained by the limited conservation outside of the *Laverania* clade. However, the phenotype in *P. falciparum* SPM3-depleted gametocytes and mosquito stages could not be further assessed in this study due to the intrinsic deficiency of the 3D7 strain to exflagellate *in vitro* and to develop in the mosquito host. Further studies with NF54 would be necessary to elucidate the role in mosquito stages but are beyond the scope of this study.

Nevertheless, to investigate the role of SPM3 in mosquito stages, we switched to the malaria mouse model. There, *P. berghei* sporozoites showed a severe defect in motility, ability to enter salivary glands, and consequently a lower transmission efficiency. This phenotype could be explained by the impaired SPMT organization due to the lack of microtubule anchoring to the IMC (Figure 5D). Microtubules in sporozoites are organized in a polar fashion with the all-but-one microtubules underlying the IMC over two thirds of the circumference and a single microtubule being located in the remaining third (61). How this asymmetry is formed is not clear but it has been speculated that this polarity contributes to the counter-clockwise continuous circular movement of *Plasmodium* sporozoites (40, 65). This suggests that displacement of microtubules from the IMC could affect persistent circular motility, as we found here for *Pb*SPM3*-*KO sporozoites (Figure 4E-G). In line, the observed helical movement is reminiscent of the motion observed for some *Plasmodium* ookinetes and *T. gondii* tachyzoites (62, 63), where the microtubules are equally spaced around the circumference.

MAPs influence several properties and define individual subpopulations of SPMT in the different stages of apicomplexan parasites. The exact role of SPM3 in microtubule organization and pellicle architecture remains to be elucidated. The close association of the SPMTs with the cytosolic face of the parasite pellicle is expected to be mediated by MAPs (11, 40, 67). We can speculate that in *P. falciparum* SPM3 in gametocytes provides specialized contact sites between SPMT and IMC that support SPMT organization, cell elongation and ultimately falciform cell shape. Consequently, loss of SPM3 may disrupt the interaction to IMC components such as PhIL1, alveolins and other IMC proteins leading to SPMT disorganization and rounding of the cell. Additional *Pf*SPM3 specific interaction partners in gametocytes remain to be identified. In accordance, our ultrastructural analysis in *P. berghei* sporozoites suggest that *Pb*SPM3 might connect the SPMTs to the IMC. Noteworthy the function of SPM3 appears to be early in gametocyte development since the *Pf*SPM3 signal vanishes prior to the disassembly of SPMTs in stage IV/V gametocytes. Association of *Pf*SPM3 to the SPMT may also influence stability and depolymerization of SPMT as shown for *T. gondii* SPM1, TLAP2, and TLAP3, MAPs that protect the stability of the SPMT in a region-dependent manner (32).

In conclusion, we identified here a *Plasmodium*-specific microtubule-associated protein with essential functions for the gametocyte and sporozoite development and hence critical for malaria transmission.

## METHODS

### *P. falciparum* culture

Blood stages of *P. falciparum* 3D7 (68) were cultured in human red blood cells (O+ or B+, Blood bank, Universitätsklinikum Hamburg-Eppendorf). Cultures were maintained at 37°C in an atmosphere of 1% O_2_, 5% CO_2_ and 94% N_2_ using RPMI complete medium containing 0.5% Albumax according to standard protocols (69). In order to obtain highly synchronous parasite cultures late schizonts were isolated by percoll gradient (70) and cultured with fresh erythrocytes for 4 hours. Afterwards sorbitol synchronization (71) was applied in order to remove remaining schizonts resulting in a highly synchronous ring stage parasite culture with a four-hour age window.

Induction of gametocytogenesis was done as previously described (54, 72). Briefly, GDV1-GFP-DD expression was achieved by addition of 2 or 4 μM Shield-1 to the culture medium and gametocyte cultures were treated with 50 mM N-acetyl-D-glucosamine (GlcNAc) for five days starting 72 hours post Shield-1 addition to eliminate asexual parasites (73). Alternatively, asexual ring stage cultures with >10 % parasitemia, cultured in the presence of choline, were synchronized with Sorbitol (71) and washed twice in choline-free RPMI medium. Cells were kept in choline-free medium for the entirety of the assay. After one reinvasion cycle cultures at trophozoite stage were treated with 50 mM N-acetyl-D-glucosamine (GlcNAc) (73) and kept on this for five days. Gametocytes were maintained in RPMI complete medium containing 0.25 % Albumax and 0.25 % sterile filtered human serum (Interstate Blood Bank, Inc. Memphis, TN, USA).

### Cloning of plasmid constructs for parasite transfection

For endogenous tagging of *Pf*SPM3 (PF3D7_1327300) using the SLI system (42) and glmS based conditional knockdown (53, 74) a 1021 bp homology region was amplified using 3D7 gDNA and cloned into pSLI-PMRT1-GFP-glmS (75) using the NotI/MluI restriction site.

For SLI-based targeted gene disruption (SLI-TGD) of *Pf*SPM3 a 399 bp homology region was amplified using 3D7 gDNA and cloned into the pSLI-TGD plasmid (42) using NotI and MluI restriction sites.

For overexpression constructs the full length sequences of PhIL1 (PF3D7_0109000), SPM1 (PF3D7_0909500) and PF3D7_1345600 were amplified from parasite cDNA and cloned into pARL-^*ama1*^ AIP-mCherry-yDHODH (41) using the XhoI/KpnI restriction site. Oligonucleotides used to generate the DNA fragments are summarized in Table S1.

For co-localization experiments the plasmids pARL-^*ama1*^ ALV5-mCherry (41), pARL-^*ama1*^ARO-mCherry (41) were used.

For gene deletion of *Pb*SPM3 (PBANKA_1342500), both a 3’ and a 5’ homology region (750 and 717 bps) were amplified from PbANKA wild-type-genomic DNA and cloned into the Pb262 vector (76) using HindIII/XhoI and EcoRI/EcoRV restriction sites. The Pb262 vector contains the *hDHFR* gene that allows for positive selection using the drug pyrimethamine. Prior to transfection, the vector was linearized using SacII and PmeI restriction enzymes followed by ethanol-precipitation.For gene deletion of *Pb*SPM3 (PBANKA_1342500), both a 3’ and a 5’homology region (750 and 717 bp) were amplified from PbANKA wild-type-genomic DNA and cloned into the Pb262 vector (76) using HindIII/XhoI and EcoRI/EcoRV restriction sites. The Pb262 vector contains the *hDHFR* gene that allows for positive selection using the drug pyrimethamine. Prior to transfection, the vector was linearized using SacII and PmeI restriction enzymes followed by ethanol-precipitation.

For endogenous tagging of *Pb*SPM3 (PBANKA_1342500), a 992 bp long 3’ homology region was amplified from PbANKA wild-type genomic DNA. The reverse primer encoded the same linker sequence that was used for GFP-tagging of *Pf*SPM3 as described in Birnbaum *et al*. 2017 (42). The amplicon was cloned into the pL18 vector (77) using EcoRI/BamHI restriction sites. Prior to transfection, the vector was linearized using the SwaI restriction enzyme followed by ethanol-precipitation.

All oligonucleotides used to generate DNA fragments as well as those used for genotyping PCRs are listed in table S1.

### Transfection of *P. falciparum*

For transfection, Percoll-purified (70) parasites at late schizont stage were transfected with 50 μg plasmid DNA using Amaxa Nucleofector 2b (Lonza, Switzerland) as previously described (78). Transfectants were selected using either 4 nM WR99210 (Jacobus Pharmaceuticals), 0.9 μM DSM1 (79) (BEI Resources). In order to select for parasites carrying the genomic modification via the SLI system (42), G418 (ThermoFisher, USA) at a final concentration of 400 μg/mL was added to a culture with about 5% parasitemia. The selection process and integration test were performed as previously described (42).

### Generation of *P. berghei* parasite lines

The linearized *Pb*SPM3-KO vector was transfected into an unmodified *P. berghei* strain ANKA using standard protocols (80). Parasites that integrated the desired DNA construct were selected by administration of pyrimethamine (0.07 mg/ml) via the mouse drinking water. Parasites from transfections are mixed populations and hence a limiting dilution was performed to isolate clonal lines. For this a single blood stage parasite was injected into each of 8 Swiss mice intravenously. Once mice reached between 1–3% parasitemia, blood was collected via cardiac puncture from anesthetized mice (100 mg/kg ketamine and 3 mg/kg xylazine, Sigma-Aldrich). Whole blood aliquots were stored in liquid nitrogen. Genomic DNA was isolated from whole blood and tested for correct construct integration by PCR. Erythrocytes were lysed in 1.5 ml phosphate buffered saline (PBS) containing 0.03% saponin. After centrifugation and washing, the genomic DNA was isolated using the Blood and Tissue kit (Qiagen Ltd) according to the manufacturer’s protocol. All generated isogenic parasite lines were analyzed via PCR. Parasite lines with the correct genotype were assumed to be isogenic.

### Determination of *P. berghei* asexual blood stage growth rate

Blood smears from all mice used for limiting dilution and carrying an isogenic parasite line were done. The parasitemia determined on day 8 post infection was used to back-calculate the growth-rate as described (81, 82).

### Mosquito infection

Frozen parasite stocks were thawed and injected intraperitoneally (100–150 μl) into one mouse. Parasites were allowed to grow for 4-6 days with the infection rate being monitored by blood smears. Once the infected mouse reached 2–3% parasitemia, the mouse was anesthetized (100 mg/kg ketamine and 3 mg/kg xylazine, Sigma-Aldrich) and bled by cardiac puncture. From this donor mouse, 20 million parasites were transferred into two naïve recipient mice by intraperitoneal injection. Three days after fresh blood transfer, exflagellation of parasites was checked first and mice were anesthetized (120 mg/kg ketamine and 16 mg/ml xylazine) to be then placed on a cage containing around 200 female *Anopheles stephensi* mosquitoes, which were starved for 3 to 5 hours by removal of sugar and salt pads. Mosquito were allowed to feed on mice for at least 15 min up to 30 min. Infected mosquitos were kept at 21°C and 70% humidity.

### Exflagellation assay *in vitro*

A drop of tail blood from an infected mouse was placed onto a glass-slide, tightly covered with a cover glass and placed at 21 °C for 12 min. Exflagellation centers were observed by light-microscopy at 40x magnification with a phase contrast ring at a Zeiss Axiostar microscope. The number of exflagellation centers was roughly counted at several fields with comparable erythrocyte densities.

### Analysis of oocyst development by mercurochrome staining

Midguts of infected mosquito were isolated on day 11 and day 12 post mosquito blood meal from three independent mosquito feeds. A minimum of 50 mosquitos per feed replicate were dissected. Midguts were dissected in 100 μl of PBS and permeabilized with 1% Nonidet P40^®^ in PBS (AppliChem GmbH) for 20 min at room temperature. The supernatant was removed and midguts were stained for 30 min up to two hours in 0.1% mercurochrome in PBS (NF XII, Sigma-Aldrich). The staining solution was discarded and midguts were washed several times with PBS until the supernatant became clear. Stained midguts were transferred onto a glass-slide leaving some PBS on top and sealed with a coverslip. Midguts were imaged using an epifluorescence microscope (CellObserver, Zeiss) using a 10x (NA 0.5, air) objective with a green filter and both the infection rate and the number of oocysts were determined.

### Sporozoite isolation, counting and imaging

Midgut and salivary gland sporozoites of infected mosquitoes were isolated on day 18 and day 20 post mosquito blood meal from three independent mosquito feeds. A minimum of 10 mosquitoes was dissected per count. The midguts and salivary glands were dissected on ice in RPMI and crushed with a pestle before counting using a Neubauer counting chamber. For imaging, crushed salivary glands were filled up to 1 ml with RPMI and carefully underlaid with 3 ml of 17% Accudenz. Centrifugation at 2800 rpm for 20 min at room temperature separated sporozoites from cell debris. The sporozoite containing interphase was collected (total volume of 1.4 ml) and sporozoites were pelleted for 3 min at 1,000 rpm (Thermo Fisher Scientific, Biofuge primo). The supernatant was removed and RPMI containing 3% bovine serum albumin (ROTH) was added to the sporozoites and the resulting mixtures transferred into the wells of an optical bottom 96-well plate (Thermo Fisher Scientific). The plate was centrifuged for 3 min at 1,000 rpm (Multifuge S1-R, Heraeus) and imaged using an epifluorescence microscope (CellObserver, Zeiss). Movies were obtained using a 25x (NA 0.8, water) objective at a speed of 1 frame every 3 seconds with 100 cycles imaged per movie. Movies were taken until latest 1 hour post sporozoite activation. Videos were analyzed for 5 minutes with Fiji (Version: 2.0.0 rc 64/1.51s (83). Gliding assays were carried out as technical replicates on day 19 and day 22 per cage infection. For analysis, data of two (WT) and three (*Pb*SPM3-KO) biological replicates were pooled and analysed single-blinded. Gliding motility was categorized into the different patterns as described before (84). Shortly, sporozoites motility behavior can be distinguished as following: continuous: persistently moving sporozoites for at least 50 out of 100 frames without stopping more than 10 frames in a row; partial: sporozoites moving in a circular manner but not persistently; helical: sporozoites moving in a new type of motility as illustrated in Figure 4E-F attached: sporozoites adhering to the substrate but showing no sign of movement; patch: sporozoites moving actively over a single adhesion spot in a back-and-forth manner; waver: sporozoites attached with one end and the other end moving actively through the medium; Floater: sporozoites floating in the medium without contact to the substrate.

### Mouse infection by natural transmission and intravenous injection of sporozoites

For natural transmission via mosquito bite, ten female mosquitos that had their blood meal 21 days before the experiment were transferred into cups the day before the experiment. Mosquitoes were starved for 3 up to 5 hours the next day by removal of sugar and salt pads. Naïve C57/BL6 mice were anaesthetized with 120 mg/kg ketamine and 16 mg/ml xylazine and one mouse was placed per cup allowing mosquitoes to feed for at least 15 min up to 30 min with a minimum number of 6 mosquitos to have fed. To confirm that mosquitos were infected, salivary glands of all blood-fed mosquitos were dissected as described before and sporozoite numbers were determined.

For intravenous injection of sporozoites, salivary gland sporozoites were dissected as previously described and 1000 sporozoites (in 100 μl PBS) were injected into the lateral teil vein of a naïve C57/BL6 mouse.

Daily blood smears were taken from day 3 post infection at approximately the same time as the infection experiment. Once the infected mice reached 2–3%parasitemia, they were sacrificed by cervical dislocation. Mice that did not become positive 20 days post infection were considered negative.

### Fluorescence imaging of fixed *P. berghei* sporozoites

Microtubules and SPM3 localization of fixed *Plasmodium berghei* sporozoites were visualized using a Zeiss Airyscan 2 LSM900 laser scanning confocal microscope. Both midgut and salivary gland sporozoites were isolated as described above but without subsequent Accudenz-purification. Instead, sporozoites were directly pelleted for 3 min at 13000 rpm (Thermo Fisher Scientific, Biofuge primo). The supernatant was removed and sporozoites were resuspended in RPMI containing 3% bovine serum albumin (ROTH). For imaging, 30,000 – 50,000 sporozoites were transferred into a well of a Labtek (μ-slide, 8-well ibiTreat, Ref 80826). Sporozoites were pelleted for 3 to 4 min at 800 rpm (Multifuge S1-R, Heraeus) to be then incubated for 10 min at RT allowing sporozoites to attach and start gliding while leaving the supernatant on top. Sporozoites were washed three times with RPMI before being fixed in 4% PFA in PBS at room temperature. After one hour of incubation, fixative was removed washing sporozoites three times with RPMI followed by permeabilization with 0.5% Triton X-100 in PBS for one hour at room temperature. Before incubation with primary antibody, sporozoites were again washed three times with RPMI. Sporozoites were incubated for one hour at room temperature with mouse anti-tubulin (1:500, Sigma-Aldrich, Ref T5168) to stain microtubules followed by three washing steps with RPMI. Goat anti-mouse Alexa Fluor-546 (1:500, Invitrogen, Ref A11030) was used as secondary antibody and incubated for 30 min up to one hour at room temperature before incubating sporozoites for 15 min in RPMI containing 1:1,000 diluted Hoechst-33342 from a 10 μM Hoechst-33342 stock solution (Thermo Fisher Scientific). Sporozoites were washed three times in RPMI to be subsequently imaged. Z-stacks were acquired with a z-spacing of 0.5 μm and a total of 29 layers being taken. Images were directly 3D-processed by the internal 3D-Airyscan processing option.

### Fluorescence imaging of live *P. berghei* parasites

Midguts were isolated as before and transferred onto a glass-slide leaving some PBS on top and sealed with a coverslip. Sporozoites were activated in 3% BSA in RPMI and transferred to an optical 96-well plate (Thermo Fisher Scientific). The plate was centrifuged for 3 min at 1,000 rpm (Multifuge S1-R, Heraeus) and imaged using an epifluorescence microscope (CellObserver, Zeiss). Images from both oocysts and sporozoites were obtained using a 63x (NA 1.4 oil) objective with an exposure of 80 m/s.

### Transmission electron microscopy (TEM)

Highly infected midguts were fixed by incubation in 4% paraformaldehyde and 4% glutaraldehyde diluted in 100 mM sodium cacodylate buffer at 4°C overnight. Fixed samples were washed three times in 100 mM sodium cacodylate buffer at room temperature (RT) for 5 min. A secondary fixation was done in 1% osmium (in 100 mM sodium cacodylate buffer) at RT for 60 min. Samples were washed twice with 100 mM sodium cacodylate buffer and twice in ddH2O and then contrasted with 1% uranyl acetate (in ddH_2_O) at 4°C overnight. Samples were washed twice with ddH_2_O for 10 min and then dehydrated by incubating in increasing concentrations of acetone (30%, 50%, 70%, 90%) for 10 min and two times in 100% for 10 min. Samples were adapted to ‘Spurr’ solution (23.6% epoxycyclohexylmethyl-3,4-epoxycyclohexylcarboxylate (ERL); 14.2% ERL-4206 plasticizer; 61.3% nonenylsuccinic anhydride; 0.9% dimethylethanolamine) by incubating in increasing concentrations (25%, 50%, 75%) at RT for 45 min and at 100% at 4°C overnight. MGs were resin embedded with ‘Spurr’ at 60°C overnight. Embedded midguts were trimmed and 70 nm thick sections were imaged on a transmission electron microscope at 80 kV (JEOL JEM-1400) using a TempCam F416 camera (Tietz Video and Image Processing Systems GmbH, Gautig).

Microtubule numbers and distances were measured in a single-blinded way: After having defined all sporozoites with clearly visible microtubules, the number of microtubules per sporozoite were manually counted. The distance of microtubules to either the IMC or a membrane of a so far unknown origin were measured by taking the distance from the center of the microtubule to the inner side of the IMC or to the membrane, respectively. A cut-off value of 40 nm was set with distances of less or equal to 40 nm being considered as “close” according to Ferreira *et al*. (26).

### Fluorescence imaging of *P. falciparum* infected erythrocytes

All fluorescence images were captured using a Zeiss Axioskop 2plus microscope with a Hamamatsu Digital camera (Model C4742-95) or a Leica D6B fluorescence microscope equipped with a Leica DFC9000 GT camera and a Leica Plan Apochromat 100x/1.4 oil objective.

Microscopy of live parasite-infected erythrocytes was performed as previously described (85). Briefly, parasites were incubated in standard culture medium with 1 μg/mL Hoechst-33342 (Invitrogen) for 15 minutes at 37°C prior to imaging. 5.4 μL of infected erythrocytes were added on a glass slide and covered with a cover slip. Nuclei were stained with 1 μg/mL Hoechst-33342 (Invitrogen). Microtubules were visualized by incubation of parasites in medium containing 1:1,000 TubulinTracker Deep Red (Thermo Fischer Scientific; dissolved in DMSO), which labels polymerized tubulin, for 15 min at 37°C prior to imaging, as previously described (86). Images were processed using Fiji (83) and Adobe Photoshop CC 2021 was used for display purposes only.

### *P. falciparum* blood stage growth assay

For growth assays of TGD cell lines a flow cytometry assay, adapted from previously published assays (87, 88), was performed to measure proliferation over five days. Each day parasite cultures were resuspended and 20 μL samples were transferred to an Eppendorf tube. 80 μL RPMI containing Hoechst-33342 and dihydroethidium (DHE) was added to obtain final concentrations of 5 μg/mL and 4.5 μg/mL, respectively. Samples were incubated for 20 min (protected from UV light) at room temperature, and parasitemia was determined using an ACEA NovoCyte flow cytometer.

### *P. falciparum* gametocyte quantification assay

Giemsa-stained blood smears at day 10 post induction of GDV1 expression were obtained and at least 10 fields of view were recorded using a 63x objective per treatment and time point. Erythrocyte numbers were then determined using the automated Parasitemia software (http://www.gburri.org/parasitemia/) while the number and morphology of gametocytes was determined manually in 1096–2632 (mean = 1793635) (3D7-iGP-SPM3-GFP-glmS and 3D7-iGP-SPM3-GFP-glmSM9) or 925–2742 (mean = 1711) (3D7-iGP control and 3D7-iGP-SPM3-TGD) erythrocytes per sample. The counting was performed blinded.

### *P. falciparum* GlmS-based gene knockdown

GlmS-based knockdown assay was adapted from previously published assays (53, 89). To induce knockdown, highly synchronous early ring stage parasites were split into two dishes, 2.5 mM glucosamine (GLCN) was added to one of them and parasite growth was measured by FACS after two and four parasite replication cycles. Parasite cultures were inspected daily by Giemsa smears and, if necessary, diluted to avoid growth bias caused by high parasitemia. As an additional control, the same amount of glucosamine was also added to 3D7 wild-type parasites. For all analyses, medium was changed daily, and fresh glucosamine was added every day.

GlmS-based knockdown in gametocytes was performed as previously described (90). Briefly, synchronized ring stage cultures were induced by the addition of Shield-1, as described above. At day 3 post induction the culture was spilt into two dishes and one dish was cultured in the presence of 2.5 mM glucosamine for the remaining ten days.

### Phylogenetic analysis

For phylogenetic analysis, we identified sequences of potentially orthologous genes in other species by using protein-to-protein BLAST search (blastp, default parameters, (91)) with the *Pf*SPM3 protein sequence as a query (accession XP_001350024.1). The blastp search was performed on the nr database but excluding, *Plasmodium falciparum‘* as a species. Selected representative sequences (PRCDC_1326300, PGABG01_1325400, PVX_116515, AK88_00350, PcyM_1231000, PKNH_1201300, PCOAH_00043100, PCHAS_1347100, C922_01771, PVVCY_1304510, PBANKA_1342500, PY17X_1347200, PocGH01_12035100, PmUG01_12037200, PRELSG_1200700, PGAL8A_00239500, HEP_00010500) were aligned with the NCBI tool Cobalt multiple alignment, and a phylogenetic gene tree (fast minimum evolution method based on Grishin protein sequence distances) was built via the NCBI web tools. Mega X (92) was used for visualization of the gene tree.

### Software

Statistical analyses were performed with GraphPad Prism version 6 or 8 (GraphPad Software, USA) or RStudio Team (2022, RStudio: Integrated Development Environment for R. Rstudio, PBC, Boston, MA URL http://www.rstudio.com/). Plasmids and oligonucleotides were designed using ApE (93) or SnapGene Software Version 3.2.1 (Insightful Science, available at snapgene.com).

### Animal work and ethics statements

For *in vivo* experiments, 6-8-week-old female Swiss mice obtained from Janvier labs were used. These experiments included parasite propagation and mosquito infections. For transmission experiments used to determine parasite infectivity, 4-6-week-old female C57/BL6 mice from Charles River laboratories were used. All animal experiments were performed under the authorization number G111/20 according to the FELASA-B guidelines and were approved by the Regierungspräsidium Karlsruhe.

## Acknowledgements

We thank Till Voss for 3D7-iGP parasites, Tobias Spielmann for SLI-TGD plasmid, Jacobus Pharmaceuticals for WR99210, Greg Burri for the parasitemia software, Miriam Reinig and members of the Frischknecht lab for raising mosquitoes, Marek Cyrklaff, Charlotta Funaya and Sebastian Weber for help with electron microscopy and Ulrich Schwarz, Josie Ferreira and Benjamin Liffner for helpful discussions. We acknowledge the microscopy support from the Infectious Diseases Imaging Platform (IDIP) at the Center for Integrative Infectious Disease Research and the Electron Microscopy Core Facility (EMCF) of Heidelberg University, as well as the Advanced Light and Fluorescence Microscopy (ALFM) facility at the Centre for Structural Systems Biology (CSSB). DSM1 (MRA-1161) was obtained from MR4/BEI Resources, NIAID, NIH.

## Author contribution

Conceptualization: JSWM, ABA, FF, TWG

Methodology: LPD

Investigation: JSWM, AMBI, PMR, LPD, GF, SS,

Formal Analysis: TLL

Writing original manuscript: JSWM, AMBI, PMR, ABA, DW, FF, TWG

Review & Editing: JSWM, AMBI, PMR, TLL, ABA, DW, FF, TWG

Funding Acquisition: ABA, FF, TWG

Resources: TWG

Project Administration: FF, TWG

Supervision: ABA, FF, TWG

All authors read and approved the manuscript.

---

**Figure S1:**
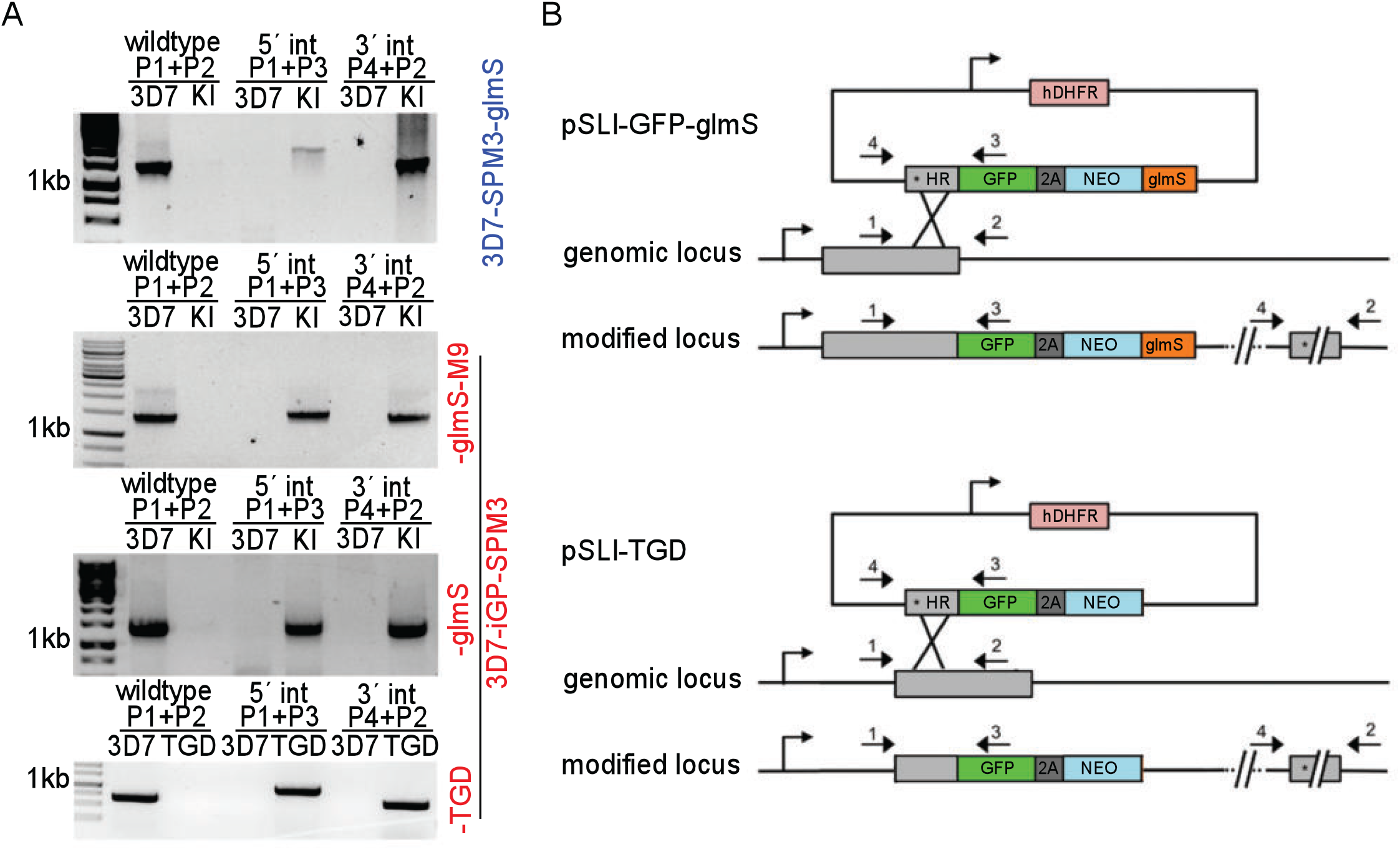
Validation of generated transgenic cell lines. **(A)** Confirmatory PCR of unmodified wild-type (WT) and transgenic knock-in (KI) cell lines to check for genomic integration at the 3’- and 5’-end of the locus. Position of the primer used are indicated with numbered arrows in the scheme. **(B)** Schematic representation of KI and TGD strategy using the selection-linked integration system (SLI) (42). Pink, human dihydrofolate dehydrogenase (hDHFR); grey, homology region (HR); green, green fluorescence protein (GFP) tag; dark grey, T2A skip peptide; blue, neomycin resistance cassette; orange, glmS sequence. Stars indicate stop codons, and arrows depict primers (P1 to P4) used for the integration check PCR.

**Figure S2:**
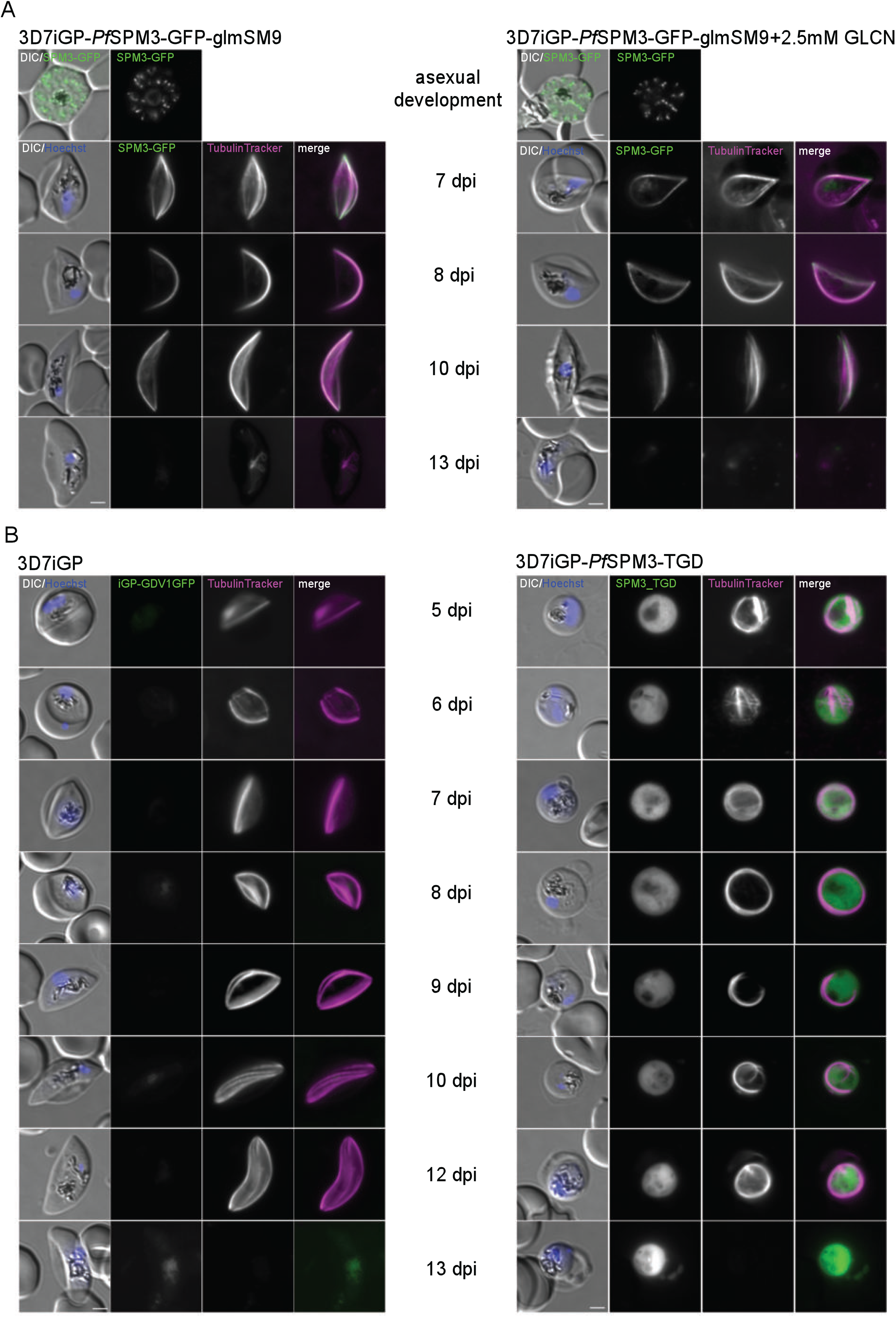
3D7-iGP-*Pf*SPM3-glmSM9 during gametocyte development. **(A)** Live-cell microscopy of 3D7-iGP-*Pf*SPM3-GFP-glmSM9 schizonts and stage II–V gametocytes (day 7 – day 13 post gametocyte induction) cultured either with or without (control) 2.5 mM GLCN. Nuclei were stained with Hoechst-33342. Scale bar, 2 μm. **(B)** Live-cell microscopy of 3D7-iGP (control) vs. 3D7-iGP-*Pf*SPM3-TGD gametocytes (day 5 – day 13 post gametocyte induction). Microtubules were visualized with TubulinTracker and nuclei were stained with Hoechst-33342. Scale bar, 2 μm.

**Figure S3:**
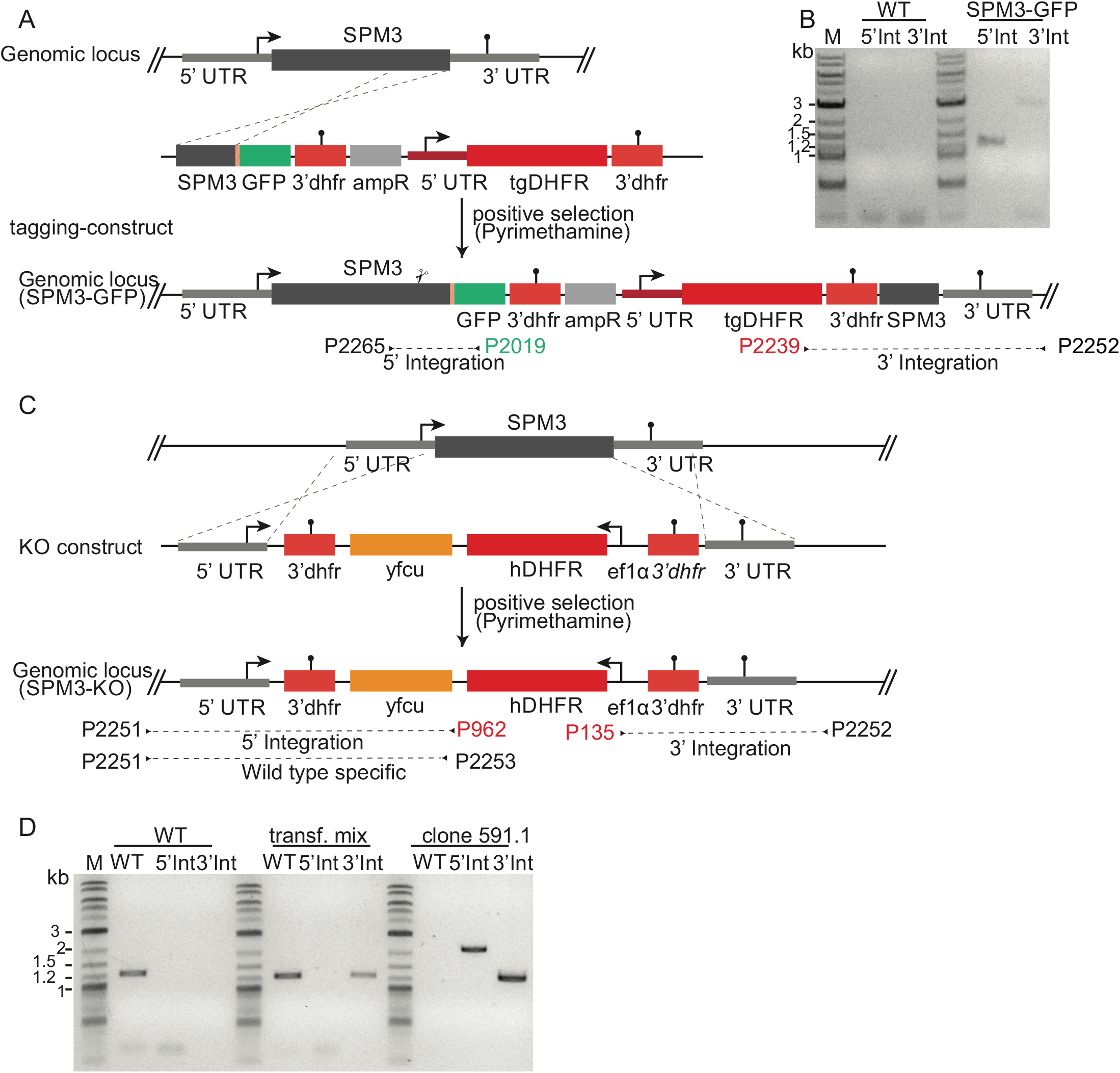
Generation of transgenic *P. berghei* parasite lines and validation. **(A, C)** Schematic representation of KI and KO strategy used to generate *Pb*SPM3-GFP parasites by single-crossover recombination (A) or *Pb*SPM3-KO parasites by double-homologous recombination (B). 3’dhfr, 3’UTR of dihydrofolate reductase; ampR, ampicillin resistance; ef1α, elongation factor 1 alpha; GFP, green fluorescent protein; tgDHFR, *Toxoplasma gondii* dihydrofolate reductase; UTR, untranslated region; yfcu, yeast cytosine deaminase and uridyl phosphoribosyl transferase. Primers are displayed in the same colour as the element they bind to (i. e. P2019 binding to *gfp*). Note that primers in black can bind to both the original as well as the mutated gene locus. **(B, D)** Genotyping PCR of the generated *Pb*SPM3-GFP (B) and *Pb*SPM3-KO (D) line in comparison to PbANKA wild-type line using primers for 3’ and 5’ integration. Binding sites of the primers used are indicated in (A).

**Figure S4:**
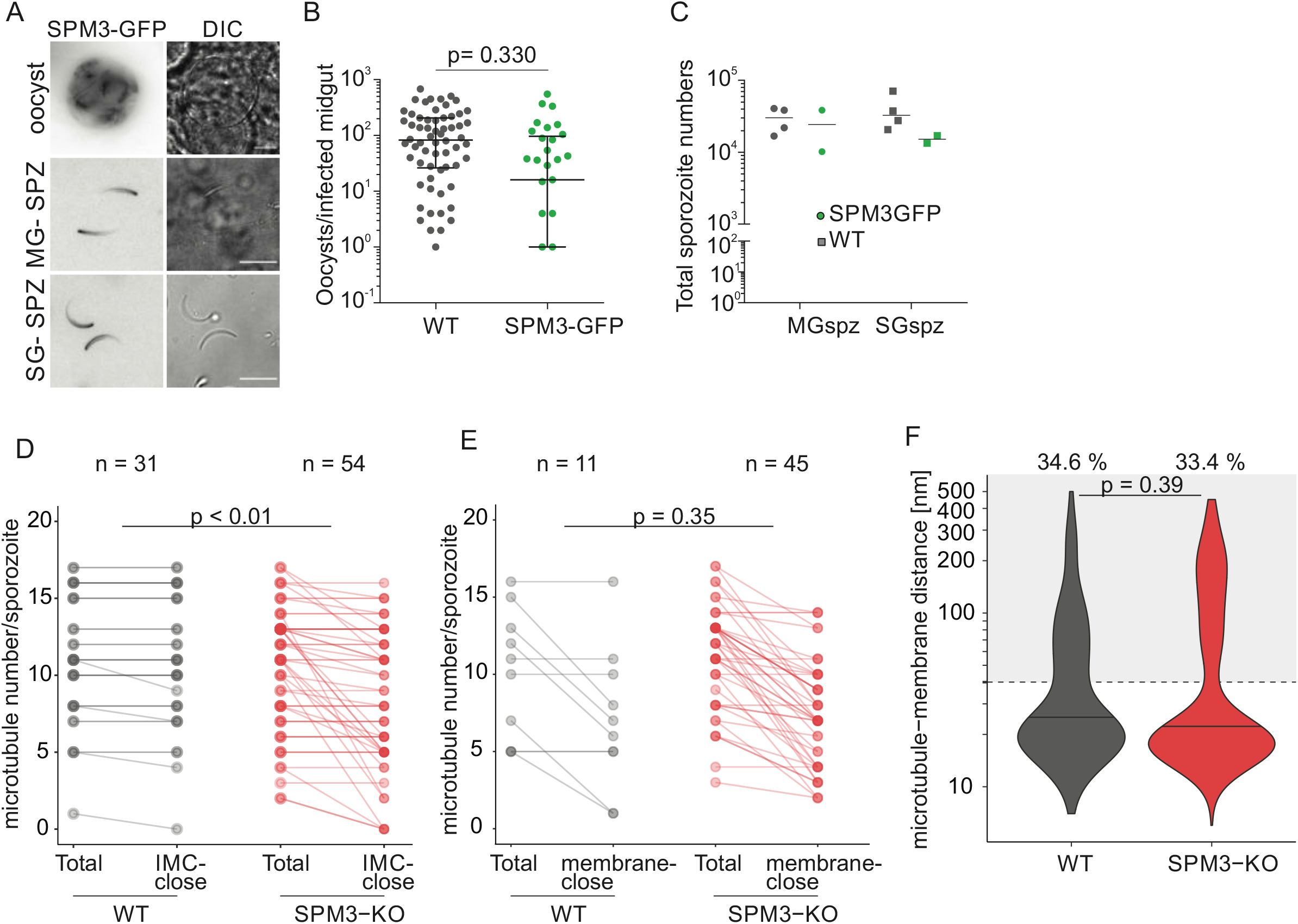
*Pb*SPM3-GFP localization and development of tagged parasite line. **(A)** Live-cell imaging of endogenously tagged *Pb*SPM3-GFP at the oocyst stage, free midgut (MG-SPZ) and salivary gland sporozoites (SG-SPZ). Scale bar 10 μm. **(B, C)** Endogenous protein-tagging has no effect on general parasite development: (B) Similar oocyst numbers of wild-type (WT) and *Pb*SPM3-KO parasites analyzed on day 11 and day 12 post infection. Overall, 78.5% (62/79) of mosquito midgut were infected with wild-type parasites compared to 66.7% (22/33) with *Pb*SPM3-GFP parasites (p = 0.23, Fisher’s exact test). For statistical analysis comparing oocyst numbers per infected midgut, Mann-Whitney test was performed. Black line indicates median with error bars representing interquartile range. **(C)** Total number of salivary gland resident versus midgut resident sporozoites determined at day 18 and 20 post infection. Black line indicates median. Note that wild-type data in B and C is used for comparison reasons. It represents the same wild-type dataset as shown in Figure 4C and D. Mosquitoes from two independent wild-type and one *Pb*SPM3-GFP replicate cage infection were analyzed. **(D)** Knockout of SPM3 leads to dissociation of microtubules from the IMC in midgut sporozoites. Absolute numbers of IMC-close (distance of ≤ 40 nm) in comparison to total microtubule numbers. In total, 31 wild-type sporozoites versus 54 *Pb*SPM3-KO-sporozoites were analyzed. The differences in opacity correspond to the number of sporozoites observed with the given microtubule number. Datapoints with higher opacity correspond to values observed in more sporozoites. Statistical difference in microtubule numbers was determined by a two-sided t-test comparing differences in numbers of IMC-close versus total microtubule numbers between the two parasite lines. **(E)** Association of microtubules to membranes in midgut sporozoites. Absolute numbers of membrane-close (distance of ≤ 40 nm) in comparison to total microtubule numbers. A total number of 11 wild-type sporozoites versus 45 *Pb*SPM3-KO-sporozoites was analyzed. The differences in opacity correspond to the number of sporozoites observed with the given microtubule number. Datapoints with higher opacity correspond to values observed in more sporozoites. Statistical difference in microtubule numbers was determined by a two-sided t-test comparing differences in numbers of membrane-close versus total microtubule numbers between the two parasite lines. **(F)** Distance between microtubules and membrane in midgut sporozoites with black line indicating median. The dashed line indicates the present cut-off value of 40 nm distance as distances equal or below 40 nm were defined as membrane-close. Grey background highlights all values being not considered membrane-close with the corresponding percentages above. A total number of 107 wild-type and 503 *Pb*SPM3-KO microtubules was measured. Spread of microtubule-membrane distances was statistically analyzed using a linear mixed model.

**Table S1: Oligonucleotides used for cloning and diagnostic genotyping PCR**

